# Caste-biased movements by termites in isolation

**DOI:** 10.1101/239475

**Authors:** Shimoji Hiroyuki, Mizumoto Nobuaki, Oguchi Kohei, Dobata Shigeto

## Abstract

The caste system of termites is an example of phenotypic plasticity. The castes differ not only in morphology and physiology, but also in behavior. As most of their behaviors within colonies involve nestmates, it is difficult to extract innate differences among castes. In this study, we focused on movement patterns of isolated individuals of *Hodotermopsis sjostedti*. We observed distinct clusters in movement patterns over 30 min, which indicates that termites have multiple innate modes of movement. The use of these modes is biased among castes, among which neotenics had a caste-specific mode and soldiers moved more actively than workers or neotenics. These caste biases may reflect different adaptive responses to social isolation. Our study provides a basis for a deeper understanding of the roles of individual movements in social behaviors.

**Summary Statement:** Movement patterns of termites in isolation were described for different castes. We proposed movements as a novel caste-specific characteristics in social insects.

## Introduction

Phenotypic plasticity refers to a flexible, epigenetic regulation of organismal phenotypes in response to environment (West-Eberhard, 2003). Because the regulation is usually adaptive to environment, phenotypic plasticity is usually considered as a product of evolutionary adaptation (Pigliucci, 2005; Pfennig et al., 2010). Phenotypes include not only morphology and physiology, but also behaviors (Hau and Goymann, 2015). While morphological plasticity is associated with ontogeny (such as molting), behavioral plasticity can emerge among individuals even with the same morphology or developmental stages. Therefore, the degree of behavioral plasticity can be characterized as crosstalk between internal (e.g., morphological, physiological, neuronal) and external (i.e., environmental, social) factors of individuals (Nussey et al., 2007; Dingemanse et al., 2009).

Insects provide plenty of examples of phenotypic plasticity, many of which are realized in response to hormones (Gilbert and Epel, 2009). Some eusocial insects, such as ants and termites, have morphologically distinct castes (Bourke and Franks, 1995; Eggleton, 2010). Each caste is a specialist of a particular colonial task associated with its morphology, and this specialization can lead to sophisticated division of labor among nestmates (Wilson, 1971). For example, soldier castes in termites have specialized mandibles that play a key role in nest defense (Eggleton, 2010). Workers of the leaf-cutting ant *Atta cephalotes* can be divided into small- and large-sized castes called minim and media. Minim ants deposit pheromones on their foraging trails more frequently than do media ants (Evison et al., 2008). Accumulating evidence indicates that social interactions among nestmates alter individual hormone levels that regulate molecular mechanisms involved in caste differentiation, thus optimizing caste ratios in colonies (Fewell and Gadau, 2009; Bourke, 2011). The degree of behavioral plasticity can depend on differences in social context among workers (Tanner, 2008; Tanner and Adler, 2009) and among castes (Ishikawa and Miura, 2012; Sun et al., 2013). To understand the role of behavioral plasticity in the division of labor in social insects, it is important to reveal innate properties of individual behavior and between-caste differences, ideally in the absence of social interactions.

One of the most elemental behavioral components of animals is their movement pattern. Although simply defined as a change in the spatial location of individuals with time, movement plays a central role in determining the fate of individuals (Nathan et al., 2008). In social insects, interactions among nestmates are essential to their lives. This need may result in the evolution of efficient movement characteristics of individual social insects. For example, workers of the termite *Cornitermes cumulans* show Lévy walk characteristics in their free walking patterns, which is known to be an efficient search movement (Miramontes et al., 2014). Since each morphological caste is assigned a role in the colony, we can hypothesize that these roles are correlated with caste-specific movement patterns. However, caste-specific patterns are largely unexplored.

The Japanese damp-wood termite *Hodotermopsis sjostedti* is a relatively basal species within termite phylogeny (Legendre et al., 2013). The elder instar larvae, called pseudergates, have the potential to differentiate into alates, neotenics (supplementary reproductives), and soldiers (Miura, 2001). Because the fourth instar and older larvae behave as the worker caste (Shimoji et al., 2017), we use the term “worker” instead of “elder instar larva” in this paper. This species provides a good model system to study caste differentiation, as morphological caste differentiation is regulated by nestmate interactions that induce hormonal changes in individuals (Cornette et al., 2008; Shimoji et al., 2017).

In this study, we tracked free-walking behavior of termites, and quantitatively evaluated the innate properties of individual movement using a video tracking system. From video analysis, we extracted movement characteristics of workers, soldiers, and neotenics. We also analyzed 14 morphological traits. By comparing innate movement properties with morphological traits, we explored the potential of movement patterns as caste-specific identifiers. We discuss how variation in innate movement patterns among castes operates in termite society.

## Materials and methods

### Termites

We collected colonies of *Hodotermopsis sjostedti* on Amami Island, Kagoshima prefecture, in April 2017. All colonies were brought back to the laboratory within their substrates, and were maintained at room temperature in the laboratory until the experiments. Sex was determined from external morphology (Miura et al., 2000). We chose females and males of three castes: workers, neotenics, and soldiers. The soldiers were easily identified by their large pigmented heads, and the neotenics were identified later by distinctly developed gonads found in their dissected abdomens (Oguchi et al., 2016).

### Experimental design

To compare the movement patterns of individuals of each caste (Fig. 1A), we recorded individual walking trajectories in an experimental arena (Fig. 1B). Each individual was marked with a black spot on the abdomen to make it easy to see. A single termite was placed in the arena and its movement was recorded for 35 min. We made the arena which was consisted of a white circular polystyrene surface (290-mm inside diameter) bounded by a plastic wall (50-mm height). The surface was cleaned with 70% ethanol before each trial. The arena was placed in a cardboard box to exclude any natural light and air currents, and the experiment was conducted at constant 25 ± 1 °C under infrared light from a LED. A web camera (DC-NCR300U, Hanwha Japan) was positioned perpendicularly above the arena so that the arena filled the image frame (Fig. 1C). Video was recorded to a Windows PC in CCI-Pro-MR software (http://www.cosmosoft.org/CCI-Pro-MR/) at a resolution of 640 × 480 pixels and a frequency of 30 FPS. The coordinates of termite movement with time were extracted from each video in UMA tracker software (http://ymnk13.github.io/UMATracker/). After each recording, each individual was stored in 70% ethanol to confirm sex and caste under a microscope. In total, we recorded the movements of 10 male workers (3 from colony A, 4 from B, 3 from C), 10 female workers (3, 4, 3), 8 male soldiers (3, 2, 3), 9 female soldiers (3, 3, 3), 3 male neotenics (2, 0, 1), and 9 female neotenics (3, 3, 3) (Table S1).

**Figure 1.**
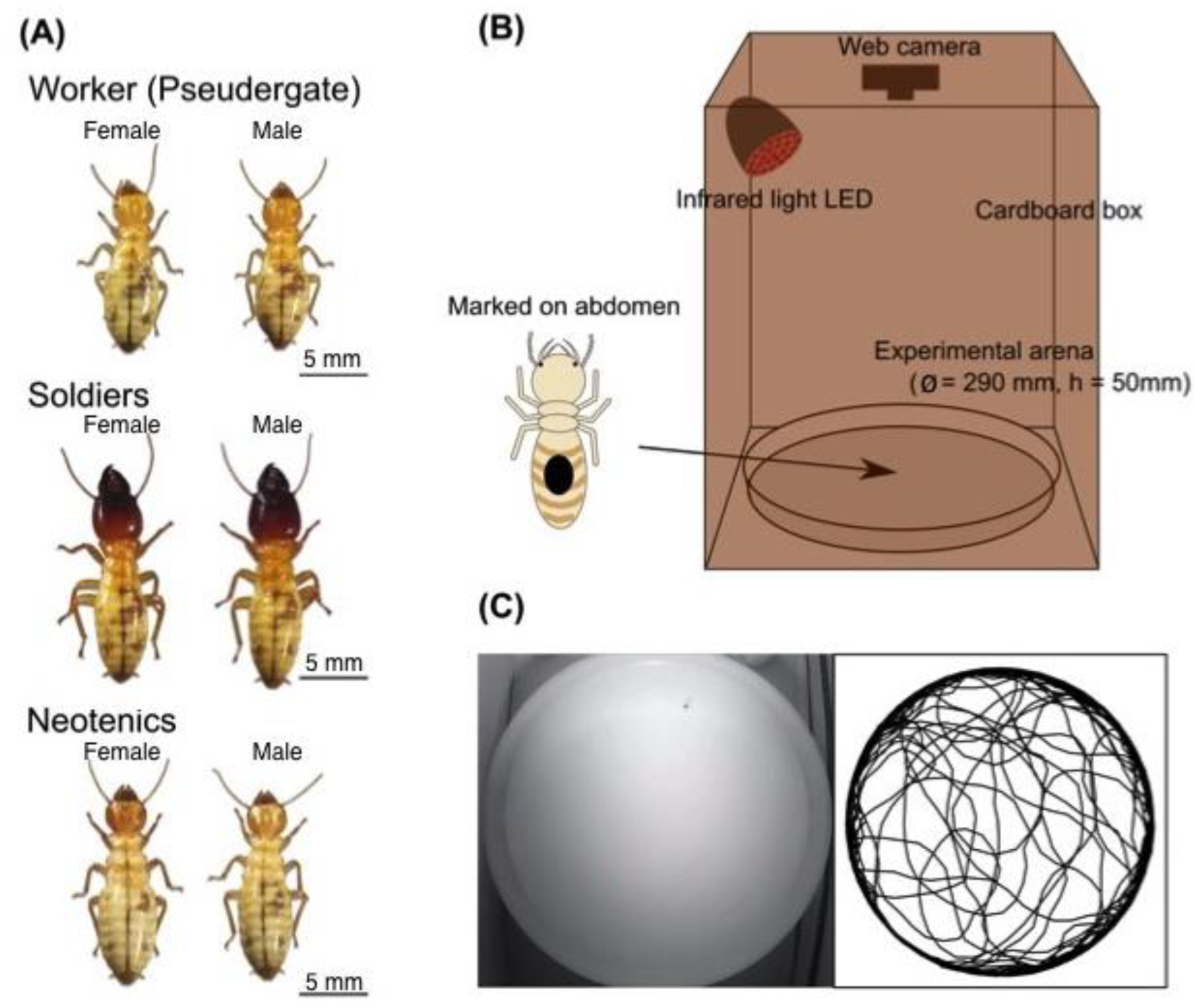
Experimental procedure. (A) Castes and sexes used. Soldiers are easily identified by their large pigmented heads. Externally, neotenics are similar to workers, so they were confirmed by dissecting the abdomen after the experiment. (B) Observation equipment: an experimental arena in a cardboard box, recorded under infrared light. (C) A video capture and an example of behavioral trajectory.

### Movement analysis

Data were analyzed in R v. 3.4.1 software (R Core Team, 2017) with the packages “CircStats”, “rcompanion”, and “MASS”. We measured nine characteristics of movement patterns from the videos: mean, maximum, and mode of instantaneous speed (mm s^−1^); SD of acceleration; total pausing time (s), number of pauses (s^−1^), total wall-following time (s); and shape (scale factor ρ) and peak position (the absolute value of peak position μ) of the distribution of turning angle when it was fitted to a wrapped Cauchy distribution. To decrease the noise arising from video analysis during data analysis, we reduced the frame frequency to 2 FPS. We discarded data of the first 5 min to avoid effects of handling. Two trials had a tracking time of <35 min. As termites sometimes stumbled and overturned, we watched each video and omitted such events from analyses.

First, we computed the instantaneous speed as the distance covered by an individual from one frame to the next. As the distribution of instantaneous speed was bimodal, with peaks at around 0 and 20 mm s^−1^, we evaluated the mode of instantaneous speed during moving from the values of the second peak. Acceleration was computed as the change in the instantaneous speed. As the mean of acceleration must be 0, we calculated the SD of the acceleration as a descriptive parameter. We assumed that a termite remained motionless when the instantaneous speed was <2 mm s^−1^. As the duration of analysis differed among individuals, we standardized the sum of pausing time and the number of pauses by dividing by the duration of analyzed time. We considered that an individual showed wall-following behavior when it was <10 mm from the wall, and the sum of wall-following time was standardized as for pausing time. The direction of movement was computed as the angle between the corresponding displacement and the horizontal, and turning angle was identified as the magnitude of change in the direction of movement from one frame to the next. To identify the degree of angular correlation in termite movements, which naturally comes from local scanning behavior of animals (Bartumeus and Levin, 2008), we fitted the data of turning angle to a wrapped Cauchy distribution using the maximum likelihood estimation method. The estimates of scale factor ρ and the absolute value of peak position μ of the distribution were obtained using the *wrpcauchy.ml* function.

### Morphometric analysis

To evaluate morphological differences among castes and sexes, we measured body parts (Koshikawa et al., 2002, 2005) of all stored individuals in image analysis software (cellSens Standard; Olympus, Japan) (Fig. S1). We measured the following 14 distances under a stereomicroscope (SZX-16; Olympus, Japan): head length (from the base of the mandible to the posterior margin of the head), maximum head width, head width at base of mandibles, labrum width, post-mentum length, post-mentum width, left mandible length (straight cross length from the condyle to the tip), pronotum length, pronotum width, mesonotum width, metanotum width, femur width (hind femur), femur length (hind femur), and tibia length (hind tibia) (Fig. S1).

### Statistical analyses

To distinguish the movement and morphological patterns among castes, we used cluster analysis with Euclidean distance matrices for degree of similarity and Ward’s method for clustering. We created separate dendrograms for movement patterns (using the nine characteristics) and morphological patterns (using the 14 body part measurements). All variables were standardized by Z-transformation before analysis.

To compare dominant movement components among castes, we performed a principal components analysis (PCA) using the function *prcomp*. PCA was performed for movement patterns and morphological patterns independently, and variables were prepared as in the cluster analysis. The Kaiser–Meyer–Olkin measure of sampling adequacy gave a value of 0.6412 for the dataset of movement patterns (classified as “mediocre”) and 0.9431 for that of morphological patterns (classified as “meritorious”; Kaiser, 1974). We reduced the variables to two principal components (PC1 and PC2) in each analysis as representative characteristics. Then we compared these PC scores between castes and colonies using Scheirer–Ray–Hare (SRH) tests (Sokal and Rohlf, 1995) with the function *scheirerRayHare*. We also compared PC1 between sexes and colonies of each caste using SRH tests.

Finally, we examined the relationship between movement traits (PC1) and morphological traits (PC1). We tested the correlation for each caste and sex using Kendall’s coefficient of concordance, pooling original colonies in each caste and sex. The significance level α was adjusted to 0.00833 using Bonferroni’s method.

### Results and Discussion

The clustering of individual movement patterns gave a hierarchical structure: in some groups, individual trajectories were distinctly more similar to one another than to those in other groups (Figs. 2, S2). Clustering and heat map patterns revealed four clusters with the following characteristics: highly active movements with frequent wall-following (cluster 1); highly active movements with less frequent wall-following (cluster 2); mixed movement of cluster 1 and 2 (cluster 3); and little active movement with frequent pauses (cluster 4) (Figs. 2, S2). This suggests that *H. sjostedti* has different movement modes, corresponding to different clusters, which can be statistically distinguished (Fryxell et al., 2008).

**Figure 2.**
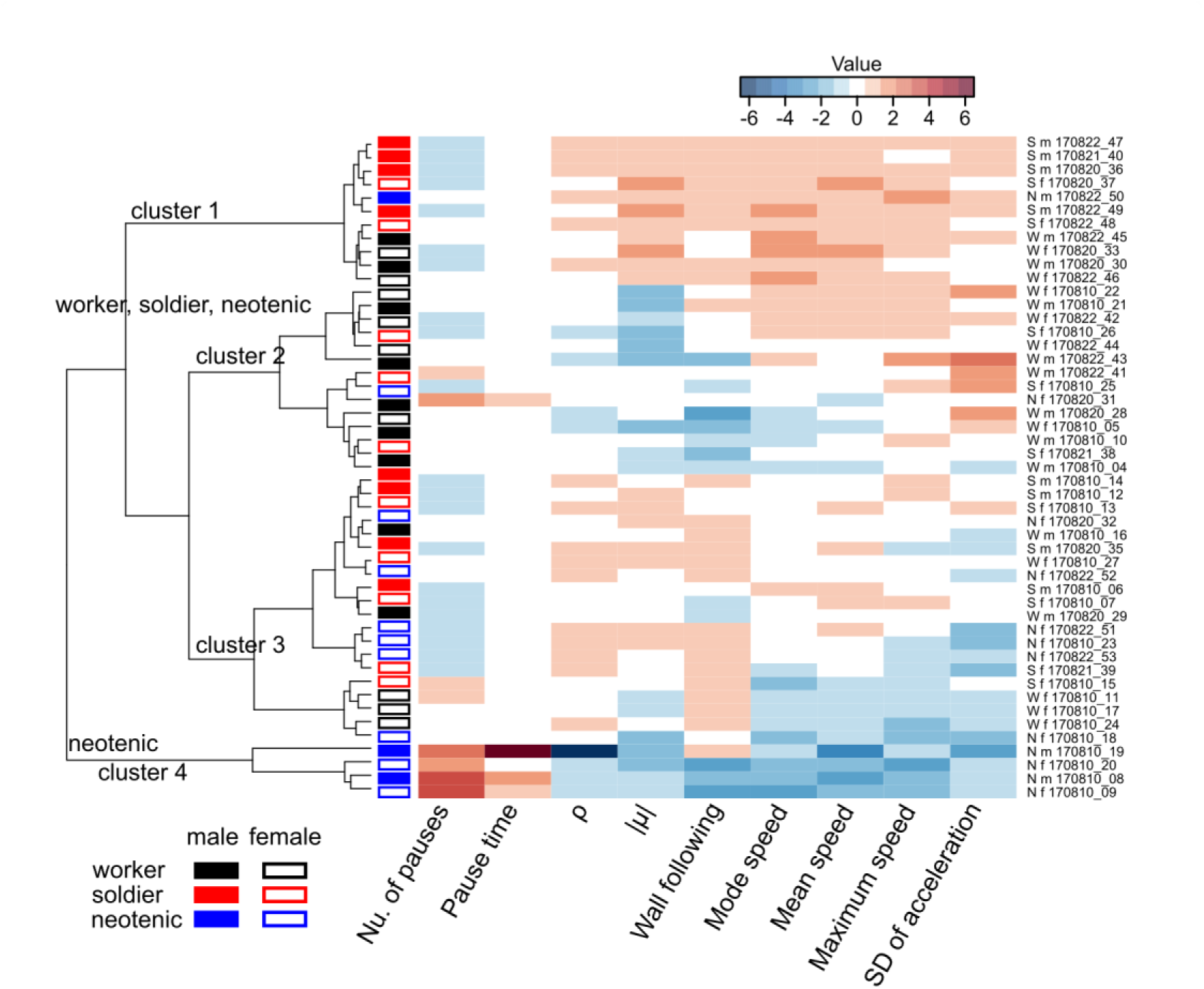
Clustering of movement patterns. Ward’s distance was calculated to make a dendrogram and to order individuals. The movement patterns were divided into four different clusters, only one of which was neotenic specific: cluster 1, highly active movement with frequent wall-following behavior; cluster 2, highly active movement with less frequent wall-following behavior; cluster 3, relatively low activity with frequent wall-following behavior; cluster 4, very low activity with frequent pauses. Values in the heat map were Z-transformed. Labels to the right indicate caste, sex, and individual: e.g., W_m_170810_4 indicates a worker male observed on 2017 August 10 with serial number 4 (see also Table S1A).

Clusters 1 to 3 encompassed all castes, whereas cluster 4 was neotenic-specific (Fig. 2). The expression of different movement modes was significantly caste-biased (Fisher’s exact test, *P* = 0.006), as soldiers favored cluster 1 and workers favored cluster 2 (Fig. 2). The first component, PC1, explained 54.51% of the total variance. The highest loadings for this component were associated with how actively termites moved (high loadings for PC1 include mean speed, pause time, and number of pauses) or moving speed (mode and maximum speed). PC2 explained 17.67% of the total variance and had the highest loadings for the total wall-following time and the SD of acceleration. PC1 was significantly different among castes and among colonies (SRH test: caste, *H*_2_ = 11.990, *P* = 0.002; colony, *H*_2_ = 10.074, *P* = 0.006; caste × colony, *H*_4_ = 3.350, *P* = 0.501; Fig. 3), whereas PC2 was not different among castes or colonies (SRH test: caste, *H*_2_ = 4.169, *P* = 0.124; colony, *H*_2_ = 0.098, *P* = 0.952; caste × colony: *H*_4_ = 6.413, *P* = 0.170; Fig. 3). A sex difference was not detected in any caste (SRH test: worker, *H*_2_ = 0.571, *P* = 0.450; soldier, *H*_2_ = 3.343, *P* = 0.068; neotenics, *H*_2_ = 0.692, *P* = 0.405).

**Figure 3.**
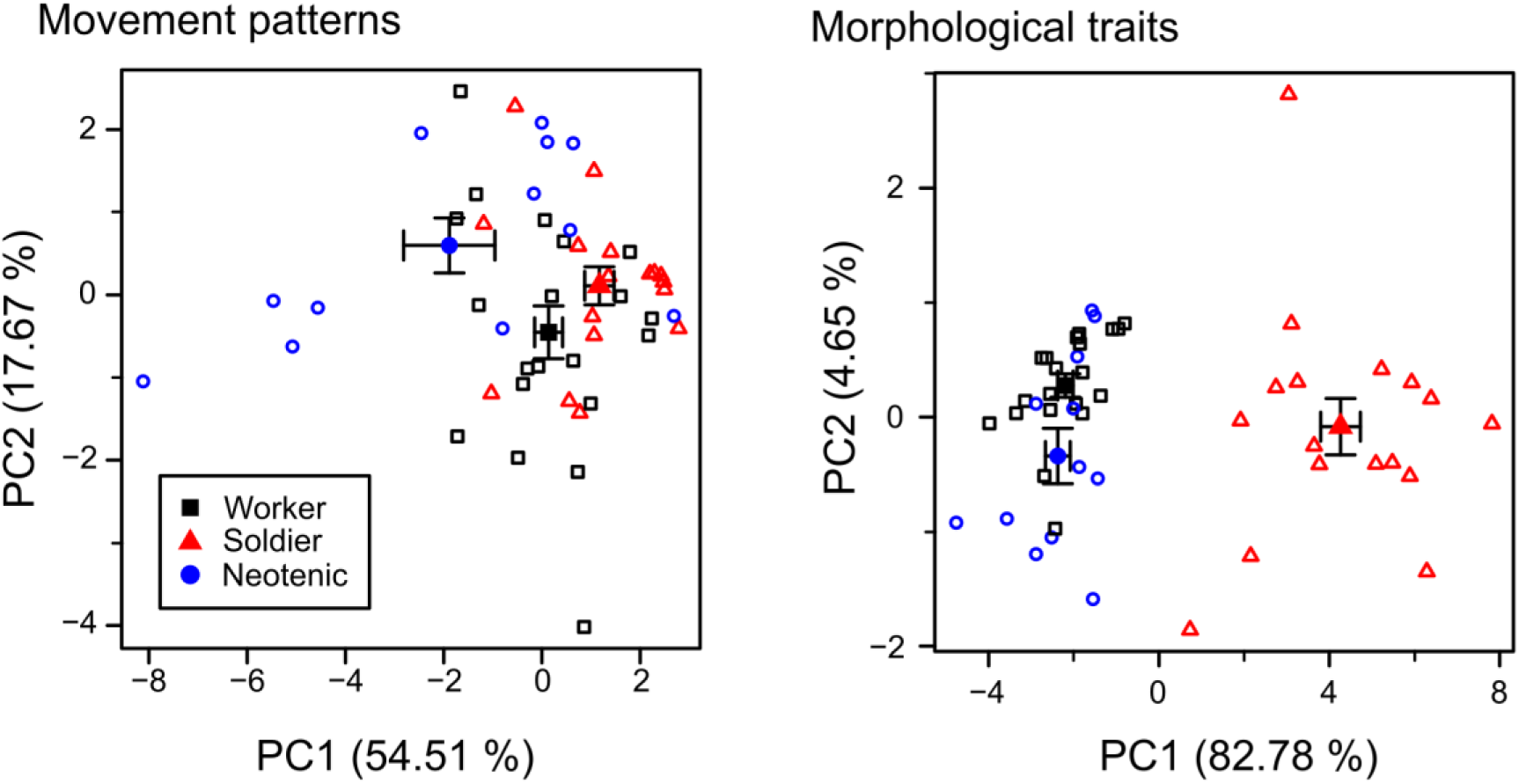
Results of principal component analysis that reduced the variables to two principal components (PC1 and PC2) for each of movement patterns and morphological patterns. □∆○ Individuals; ■▲● mean of each caste. Bars: means ± SEM.

In stark contrast, cluster analysis using individual morphology gave two distinct clusters, one of which contained all soldiers and the other was a mixture of workers and neotenics (Fig. S3). As evident from the heat map, the soldier cluster was characterized by larger size in all body parts (Figs. S3, S4B). Similarly, PCA reduced the 14 morphological characteristics to two major PCs, of which PC1 explained 82.78% of the total variance and PC2 explained 4.65%. As PC1 was positively correlated with all measurements, it was considered to indicate general body size. PC1 was significantly different among castes (SRH test: caste, *H*_2_ = 32.664, *P* < 0.001; colony, *H*_2_ = 2.558, *P* = 0.278; caste × colony: *H*_4_ = 0.821, *P* = 0.936; Fig. 3), whereas PC2 was not significantly different among castes or colonies (SRH test: caste, *H*_2_ = 5.725, *P* = 0.057; colony, *H*_2_ = 2.931, *P* = 0.231; caste × colony: *H*_4_ = 4.185, *P* = 0.382; Fig. 3). We did not find any correlation between morphological patterns (PC1) and movement patterns (PC1) in any caste or sex (female worker: *T* = 26, *P* = 0.601; male worker: *T* = 20, *P* = 0.728; female soldier: *T* = 12, *P* = 0.260; male soldier: *T* = 16, *P* = 0.720; female neotenic: *T* = 22, *P* = 0.477; male neotenic: *T* = 1, *P* = 1; Fig. S5). Overall, we concluded that the caste-biased movement patterns reflect specific characteristics of the castes themselves rather than morphological differences associated with castes.

Our results show that the emergence of different movement modes was caste biased. This indicates that movement patterns of individuals can indicate castes in termites. First, morphologically distinct soldiers showed greater activity than workers and neotenics (Figs. 3, S4A). Second, some neotenics, which cannot be distinguished from workers based on external morphological characteristics, showed a specific movement mode that was characterized by extremely low mobility with frequent pauses (Figs. 2, 3, S2). Because neotenics are the only reproductives among the castes analyzed in this study, the neotenic-specific movement mode should be related to physiological conditions involved in reproductive activity. Further investigation of individual movement patterns might enable us to find cryptic castes that cannot be distinguished in their morphology and physiology across social insects (Robinson, 2009).

As termites are social insects, the movements of isolated individuals can be regarded as searching for other nestmates (Miramontes et al., 2014). Our results show that many individuals spent a lot of time near the wall, although there was no difference among the castes (Fig. S4A). Such wall-following behavior can be seen in various social animals. For example, in ants social isolation results in increased locomotor activity and wall-following behavior (Koto et al., 2015). The wall-following behavior of woodlice results in aggregation close to the edge of the arena (Devigne et al., 2011). The same is true of schooling behavior by fish in a tank (Suzuki et al., 2003). Because physical heterogeneities affect the spatial distribution of organisms, wall-following behavior can increase the probability of encountering conspecifics. Thus, wall-following behavior might be an adaptive strategy to search for other individuals when social living animals are isolated.

In general, self-organized systems of social insects can achieve complex patterns such as the division of labor via social interactions (Camazine et al., 2001). One primary goal of research is to reveal how such complex collective behavior can emerge from individual behaviors. We found that different castes of termites in social isolation move in different ways (Figs. 2, 3), which should lead to caste-specific encounter patterns within a nest. Consequently, the rate of interactions between individuals could be determined by their movement patterns (James et al., 2008; Viswanathan et al., 2011; Mizumoto et al., 2017). The next step would be to reveal the feedback between movement patterns of individuals and encounters with other individuals in natural settings. By focusing on individual-based movement patterns as caste-specific characteristics, our study provides a novel direction for studying the division of labor in social insects.

## Acknowledgements

We thank Kenji Matsuura, Tomonari Nozaki, Yao Wu, and Ryotaro Ohi for their help with termite collection; Ryusuke Fujisawa, Naohisa Nagaya, and Masato S. Abe for fruitful discussion leading to the conception of the study; and Toru Miura for termite morphometry.

## Competing interests

The authors declare no competing interests.

## Author contributions

HS and NM designed the study and collected behavioral data. NM analyzed data. KO carried out morphometric analyses. HS, NM, KO, and SD wrote the manuscript. All authors gave final approval for publication.

## Funding

This work is supported by Grants-in-Aid for JSPS Research Fellows to NM (no. 15J02767) and KO (no. 17J06879); and for Scientific Research (B) to HS and SD (no. 15H04425).

